# A dual mechanism of sensitivity to PLK4 inhibition by RP-1664 in neuroblastoma

**DOI:** 10.1101/2025.02.15.638336

**Authors:** Isabel Soria-Bretones, Matias Casás-Selves, Elliot Goodfellow, Li Li, Cathy Caron, Ariya Shiwram, Hyeyeon Kim, Danielle Henry, Nancy Laterreur, Julian Bowlan, Alejandro Álvarez-Quilón, Frédéric Vallée, Artur Veloso, Jordan T.F. Young, Marc L. Hyer, Stephen J. Morris, C. Gary Marshall, Michal Zimmermann

## Abstract

A novel therapeutic strategy was recently proposed for high-risk neuroblastoma carrying copy number gain of the *TRIM37* gene: centriole loss upon inhibition of polo-like kinase 4 (PLK4), while tolerated by normal cells, induces aberrant mitotic spindle formation and p53-dependent cell death in *TRIM37*-overexpressing cells. Interestingly, while full PLK4 inhibition causes centriole loss, partial inhibition is known to elevate centriole numbers. Here we show using a novel selective PLK4 inhibitor RP-1664 that both centriole loss and amplification contribute to hypersensitivity of neuroblastoma cells. Whereas inactivation of *TRIM37* and *TP53* rescues neuroblastoma cell death at higher concentrations of RP-1664, at lower doses cell death is *TRIM37/TP53*-independent. With CRISPR screens and live cell imaging we demonstrate that upon centriole amplification, neuroblastoma cells succumb to multipolar mitoses due to inability to cluster or inactivate supernumerary centrosomes. *In vivo*, RP-1664 shows robust efficacy in neuroblastoma xenografts at doses consistent with centriole amplification.

**STATEMENT OF SIGNIFICANCE:** High-risk neuroblastoma is associated with poor outcomes in pediatric patients and novel therapies need to be developed. We show that neuroblastoma cells are remarkably sensitive to PLK4 inhibitors due to a combination of two complementary mechanisms, supporting the evaluation of PLK4 inhibitors in clinical trials of high-risk neuroblastoma.

## INTRODUCTION

Neuroblastoma (NBL) is the most frequent extracranial solid tumor in pediatric patients. About half of NBL cases classify as a high-risk group with poor prognosis and low overall survival rates, in particular upon relapse from standard therapy^1–4^. Novel treatment options therefore need to be developed.

Partial genomic gains at the long arm of chromosome 17 (17q) by unbalanced translocation are among the most recurrent molecular alterations in NBL^5–8^ (reviewed in^9,10^). 17q gain is observed in at least 50% of NBL cases and is an important marker of poor prognosis, with ∼70-80% of stage 4 tumors showing 17q gain^5^. Although 17q gains in NBL vary in size, they most often span a region between 17q11 to 17q25, which contains the *TRIM37* gene located in the 17q23 locus^5,11,12^. While the role of *TRIM37* gain in NBL tumorigenesis is unclear, it was proposed to confer a therapeutic vulnerability due to synthetic lethality with inhibition of polo-like kinase 4 (PLK4)^13,14^.

PLK4 is a dimeric serine-threonine kinase that plays key roles in centrosome formation^15^. The centrosome, the main microtubule-organizing center of the cell, is composed of two centrioles and the peri-centriolar material (PCM), which together nucleate microtubules that form the mitotic spindle^15,16^. The formation of centrosomes is tightly regulated. Each mitotic cell normally has precisely two centrosomes (one at each pole of the mitotic spindle), which drive bipolar division and equal separation of the mother cell’s genetic material (as well as centrosomes) into the two daughters. In turn, each G1 daughter cell will contain one centrosome with one centriole doublet that is duplicated in S-phase, ensuring that the subsequent mitosis will again contain two centrosomes^16^. PLK4 is critical for centriole duplication in S-phase: it interacts with, and phosphorylates, several centriolar and PCM proteins to drive *de novo* procentriole assembly on the parent centriole^17–21^. Consequently, PLK4 downregulation leads to progressive loss of centrioles, whereas PLK4 overexpression leads to centriole amplification and supernumerary centrosomes^17,18,22^. To guarantee that centriole duplication occurs only once per cell cycle, PLK4 also regulates its own stability: trans-autophosphorylation of several serine and threonine residues within a PLK4 dimer activates degradation of the protein by the SCF-βTrCP ubiquitin ligase^22–25^. Catalytically inactive PLK4 therefore accumulates in the cell^22–25^.

Synthetic lethality between high TRIM37 levels and PLK4 inhibition is based on their complementary roles in centrosome biogenesis^13,14^. PLK4 inhibition leads to centriole loss, for which normal cells compensate in part by using the PCM to nucleate the mitotic spindle^13,14^. However, as *TRIM37* encodes an E3-ubiquitin ligase that negatively regulates the stability of multiple PCM components^13,14,26^, high TRIM37 levels compromise the PCM’s integrity. PLK4 inhibition in the context of high TRIM37 abundance thus leads to depletion of both centrioles and the PCM, resulting in loss of centrosomes, inability to form the mitotic spindle, and mitotic failure or delay^13,14^. Prolonged mitosis is sensed by the mitotic surveillance (or ‘stopwatch’) pathway driven by the USP28-53BP1-p53 axis, which triggers a p21-mediated cell cycle arrest and cell death^13,14,27–30^. High TRIM37 levels and functional p53 therefore together drive sensitivity to PLK4 inhibition in tumor cells, whereas healthy cells with normal TRIM37 levels are more resistant due to a functional PCM^13,14,27–30^. Of note, TRIM37 has been shown to modulate sensitivity to PLK4 inhibition also by non-catalytic roles, in addition to regulating the PCM’s stability^31^. Based on the TRIM37-PLK4 synthetic lethality paradigm, we developed RP-1664: a highly selective, potent, and orally bioavailable PLK4 inhibitor (PLK4i; full characterization of RP-1664 will be reported in a separate manuscript). RP-1664 is currently being evaluated in clinical trials (NCT06232408).

The effect of pharmacological PLK4 inhibition on centriole biogenesis is bimodal. Whereas complete PLK4 inactivation at higher concentrations of PLK4i leads to centriole loss, lower concentrations have been shown to induce centriole amplification^31–34^. This centriole amplification has been attributed to an intermediate inhibition state of PLK4: enough inactive PLK4 monomers exist to reduce trans-autophosphorylation and increase total PLK4 levels, but not enough PLK4 molecules are inhibited to suppress centriole duplication^31,35,36^. Here we show that centriole amplification driven by low doses of RP-1664 contributes to PLK4i-sensitivity of NBL tumor cells. We demonstrate that many NBL cell lines are hypersensitive to RP-1664 at doses below those required for centriole loss and that sensitivity at these lower doses occurs independently of *TRIM37* and *TP53* status. Instead, NBL cells are sensitive to low doses of RP-1664 due to their inability to compensate for supernumerary centrosomes. We propose that NBL presents a uniquely suited target population for PLK4i, due to the presence of two independent mechanisms of sensitivity: one driven by centriole depletion and *TRIM37* gain at higher doses, and one driven by centriole amplification at lower doses.

## RESULTS

### RP-1664 is a PLK4 inhibitor with pre-clinical activity against TRIM37-amplified, TP53-WT tumors

RP-1664 is a novel, highly potent, selective and orally bioavailable PLK4 inhibitor (full characterization of RP-1664 will be reported in a separate manuscript). To confirm that RP-1664 inhibits PLK4 in human cells, we assessed its ability to induce PLK4 stabilization, modulate centrosome numbers, and activate the p53 mitotic surveillance pathway (**Figure 1A-D; Supplementary Figure 1A,B**). First, we analyzed the effect of RP-1664 on PLK4 protein level, as inhibition of PLK4 counteracts its SCF-βTrCP-mediated degradation^22–25^. As expected, RP-1664 treatment led to a dose-dependent increase in total PLK4 protein in RPE1-hTERT Cas9 *TP53-*null *(KO)* cells^37^ when measured by capillary-based immunodetection, with a maximal effect observed at concentrations of above 100 nM and partial PLK4 stabilization occurring below this dose (**Figure 1A,B**). To determine whether RP-1664 disrupts PLK4-dependent centriole biogenesis, we utilized a high-content fluorescence microscopy assay to quantify the number of centrosomes (visualized by anti-γ-Tubulin immunofluorescence^38^) per mitosis (marked by phosphorylated serine 10 on histone H3 – H3pS10^39^) (**Figure 1C,D**). In both p53-proficient and -deficient RPE1 cells, RP-1664 induced centrosome loss at concentrations >100 nM, with the majority of mitotic cells showing one or no centrosomes after ∼2 population doublings (**Figure 1C,D**). On the other hand, concentrations between ∼25-100nM induced supernumerary centrosomes, as expected from partial PLK4 inhibition^31^ (**Figure 1C,D**). Finally, to explore whether RP-1664 activates the mitotic surveillance pathway, we stained *TP53*-wild type (WT) RPE1 cells for the p53 transcriptional target p21 (**Supplementary Figure 1A**). RP-1664 increased p21 expression in a dose-dependent manner, at concentrations that were well in agreement with those inducing PLK4 stabilization and modulating centrosome numbers (**Supplementary Figure 1A,B**). In aggregate, these data show that RP-1664 is a *bona fide* inhibitor of PLK4 in cells that induces phenotypes consistent with published observations^31,32,34,40^.

**Figure 1.**
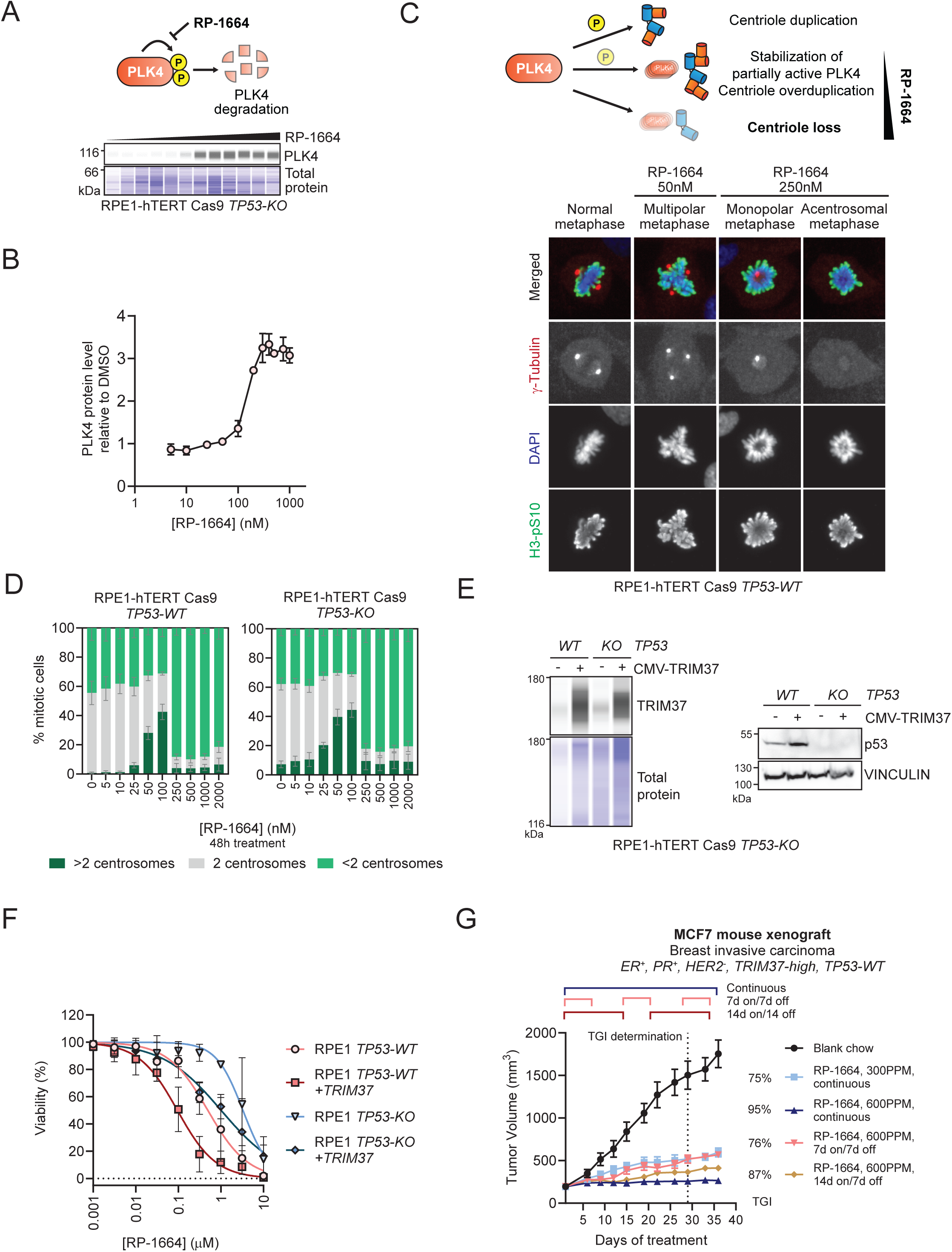
RP-1664 is a potent and selective PLK4i. A,B. RP-1664 induces PLK4 stabilization. A. Top: Model of PLK4 self-regulation. PLK4 autophosphorylation leads to its degradation. This is blocked by RP-1664. Bottom: Representative capillary immunodetection of PLK4 in RPE1-hTERT Cas9 *TP53-KO* whole cell extracts. Total protein shown as a loading control. B. Quantification of PLK4 protein levels in a representative (of *N*=3 independent experiments) capillary immunodetection assay. Mean of two technical replicates ±SD. C,D. RP-1664 modulates centrosome biogenesis. C. Top: Schematic of bimodal modulation of centriole numbers by RP-1664. Low concentrations induce centriole amplification, higher concentrations lead to centriole loss. Bottom: Representative micrographs of RPE1-hTERT Cas9 *TP53-KO* cells after no treatment of treatment with indicated RP-1664 concentrations and immunofluorescence staining with γ-Tubulin (visualizing centrosomes) and H3-pS10 (mitotic marker) antibodies. DAPI is a nuclear counterstain. D. Quantification of mitotic RPE1 Cas9 *TP53-WT* and *KO* cells with <2, 2, and >2 centrosomes at indicated RP-1664 concentrations in *N*=3 independent experiments. Mean value (bars) is shown ±SD. E-G. WT p53 and high TRIM37 sensitize to RP-1664. E. Representative (of N≥2 independent experiments) TRIM37 capillary immunodetection (left) and p53 immunoblot (right) of RPE1-hTERT Cas9 *TP53-WT* and KO cells with or without CMV-TRIM37 overexpression. Total protein and vinculin are loading controls. F. Dose-response of RP-1664 on growth (measured by Incucyte) of RPE1 *TP53-WT* and *KO* cells, with or without CMV-TRIM37. Mean of *N*=3 independent experiments ±SD. Solid lines show a non-linear regression fit to a four-parameter dose-response model. G. Tumor volume measurements of MCF7 mouse xenograft tumors in animals fed blank chow or RP-1664-containing chow at indicated doses and schedules. TGI = percent tumor growth inhibition relative to blank chow. Mean of *N*=6 mice/group ±SEM.

To evaluate the contributions of *TRIM37* and *TP53*^13,14^ to RP-1664 sensitivity, we overexpressed the TRIM37 open reading frame controlled by a CMV promoter in RPE1-hTERT Cas9 *TP53-WT* and *TP53-KO* cells using lentiviral transduction. TRIM37 transduction led to a 3-5x increase in TRIM37 protein level (**Figure 1E; Supplementary Figure 1C**). We then compared the sensitivity of TRIM37-overexpressing cells to parental *TP53-WT* and *TP53-KO* cells by IncuCyte growth assays, revealing a ∼30-fold increase in sensitivity to RP-1664 from TRIM37-normal/*TP53-KO* cells to TRIM37-overexpressing/*TP53-WT* cells, with TRIM37-high/*TP53-KO* and TRIM37-normal/*TP53-WT* cells showing intermediate sensitivity (**Figure 1F**). Importantly, concentrations of RP-1664 leading to complete growth inhibition in TRIM37-high/*TP53-WT* cells were above 100nM, consistent with the doses of RP-1664 found to induce centrosome loss (**Figure 1D**). These data confirm that high TRIM37 protein level and functional p53 cooperatively increase cellular sensitivity to PLK4 inhibition and centrosome depletion by RP-1664.

To determine the *in vivo* activity of RP-1664 against *TRIM37-high TP53-WT* tumors we engrafted the MCF7 breast carcinoma cell line subcutaneously into immunodeficient mice. MCF7 cells express estrogen and progesterone receptors but no human epithelial growth factor receptor (ER^+^, PR^+^, HER2^-^), are *TRIM37*-amplified, *TP53-WT*, and have been previously shown to be sensitive to PLK4i in a *TRIM37*-dependent manner^13^. For successful implantation and tumor growth, mice were irradiated prior to inoculation and supplemented with estradiol in their drinking water. We treated these xenograft tumors with RP-1664 delivered in mouse chow at multiple doses and schedules, with blank chow used as a vehicle control (**Figure 1G**). We observed a dose-dependent anti-tumor activity of RP-1664 with maximal tumor growth inhibition (TGI) of 95% at 600 parts per million (ppm) RP-1664 chow (**Figure 1G**). RP-1664 also demonstrated schedule flexibility, with an intermittent – two weeks on, one week off (14d on/7d off) – schedule at 600 ppm showing only modest reduction in efficacy compared to continuous delivery (**Figure 1G**). All doses and schedules were well tolerated with body weight (BW) loss of <20% in all cases, even though estradiol supplementation negatively impacted body weight loss in all groups (**Supplementary Figure 1D**). We conclude that RP-1664 shows efficacy in pre-clinical models of tumors with high levels of TRIM37 and WT p53.

### NBL cell lines are sensitive to PLK4i at doses causing centriole amplification

Due to the high prevalence of 17q / *TRIM37* gain in NBL, we sought to evaluate the pre-clinical efficacy of RP-1664 in NBL models. We assembled a panel of nine NBL cell lines carrying 17q gain and assessed their sensitivity to RP-1664 in cell viability assays (Incucyte or CellTiter Glo (CTG) as indicated in Figure legend). All cell lines showed robust sensitivity to RP-1664 with IC_50_ values in the nanomolar range (**Figure 2A**), confirming that PLK4i sensitivity of NBL models is highly penetrant.

**Figure 2.**
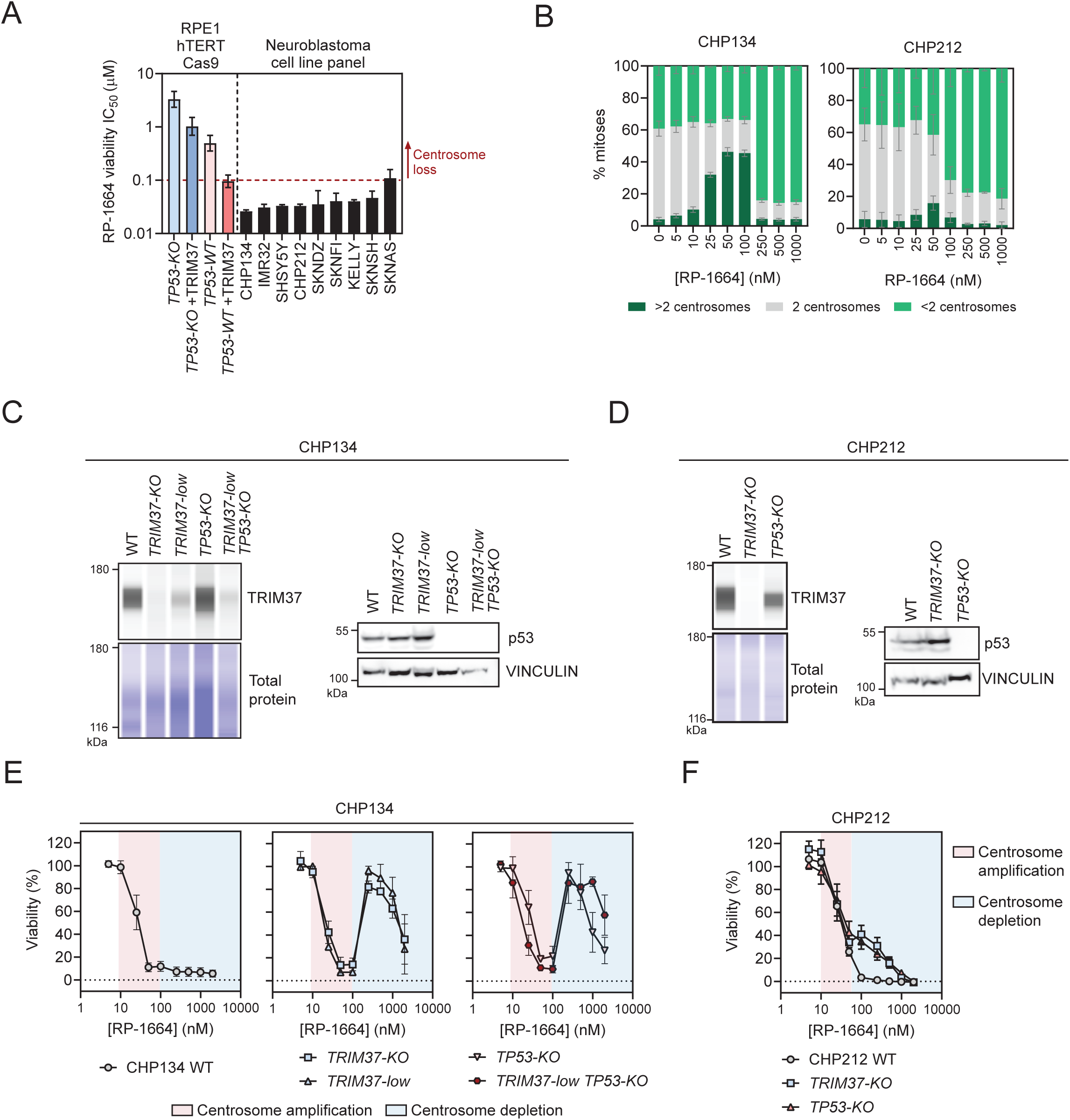
TRIM37- and p53-independent sensitivity of NBL cells to centrosome amplification. A. RP-1664 cell growth IC_50_ values for indicated cell lines. Data from non-linear least square fitting of mean viability values (N≥3 independent experiments) in growth (Incucyte (RPE1, CHP134) or CellTiter Glo (all others)) assays ±95% confidence interval. Dashed line shows the lowest concentration inducing centrosome loss in RPE1 cells. B. Quantification of mitotic CHP134 and CHP212 cells with <2, 2, and >2 centrosomes at indicated RP-1664 concentrations in *N*=3 independent experiments. Mean value (bars) is shown ±SD. C,D. Representative TRIM37 capillary immunodetection (left) and p53 immunoblots (right) of CHP134 (C) and CHP212 (D) of indicated genotypes. Total protein and vinculin are loading controls. E,F. Cell viability of CHP134 (E; measured by Incucyte) and CHP212 (F; CellTiter Glo) cells of indicated genotypes treated with indicated RP-1664 concentrations. Pink area shows concentrations causing centrosome amplification, blue represents centrosome depletion. Mean of *N*=3 to 5 independent experiments ±SD.

When evaluating the *in vitro* sensitivity of our NBL cell line panel, we noticed that the IC_50_ values of most tested cell lines fell below the 100nM concentration that is required for depletion of centrosomes in RPE1 cells (**Figures 2A, 1D**). Instead, the IC_50_ values correlated with RP-1664 doses inducing centrosome amplification. This was not due to cell type-dependent differences in modulation of centrosome number by RP-1664, as we observed centrosome amplification between 10 to ∼50-100nM, and centrosome loss at or above 100nM RP-1664 not only in RPE1 cells, but also in three different NBL cell lines (CHP134, CHP212, SHSY5Y; **Figure 2B; Supplementary Figure 2A,B**). Since it is centriole loss that underlies the sensitivity of TRIM37-high and WT p53 cells to PLK4i^13,14^, we wondered whether TRIM37 and p53 status affects the sensitivity of NBL cells to RP-1664 also at concentrations causing centriole amplification. To that end we generated *TRIM37-* and *TP53*-*KO* clones in two NBL cell backgrounds – CHP134 and CHP212 – by CRISPR/Cas9-enabled gene editing (**Figure 2C,D**). Furthermore, we derived a clone of CHP134, which we refer to as *TRIM37-low*, that retains TRIM37 expression at a reduced level, likely due to residual wild-type alleles (**Figure 2C; Supplementary Figure 2C**). Finally, we also made a CHP134 *TRIM37-low* / *TP53-KO* cell line, which has p53 inactivated on top of lowered TRIM37 levels (**Figure 2C).** We then assessed the RP-1664 sensitivity of these cell lines by Incucyte or CTG growth assays and compared it to parental WT cells (**Figure 2E,F**).

P53 inactivation, lowering TRIM37 levels, as well as ablating TRIM37 completely, rescued the CHP134 cell sensitivity to RP-1664 at concentrations ≥100nM (associated with centrosome loss), but were unable to induce resistance to RP-1664 at lower doses that lead to centrosome amplification (**Figure 2E**). Similarly, *TRIM37* or *TP53* inactivation in CHP212 cells reduced RP-1664 sensitivity at concentrations leading to centrosome depletion but had no effect at concentrations causing centrosome amplification, although in this model centriole depletion was required for complete cell killing (**Figure 2E,F**). The rescue of sensitivity to higher, but not lower, PLK4i doses by reducing TRIM37 level in CHP134 cells was reproduced with the published selective PLK4i Centrinone B^40^, ruling out compound-specific effects (**Supplementary Figure 2D-F**). Cellular sensitivity at different RP-1664 concentrations correlated well with p53 activation: whereas RP-1664 induced p21 expression in CHP134 cells at doses leading to centrosome amplification as well as depletion, lowering TRIM37 levels dampened p21 expression only at centrosome depletion concentrations (**Supplementary Figure 3A,B**). Furthermore, whereas a dose of RP-1664 causing centrosome depletion prolonged mitotic duration in CHP134 cells but not CHP134 *TRIM37-low* as expected from the TRIM37-PLK4 synthetic lethality model^14,27^, a dose causing centrosome amplification did not (**Supplementary Figure 3C,D**), again suggesting a different mechanism of action. Finally, we confirmed that sensitivity of CHP134 and CHP212 cells to RP-1664 doses causing centrosome amplification and depletion correlated with induction of cell death and apoptosis as measured by Annexin V and Cytotox (reflective of membrane permeability) staining (**Supplementary Figure 3E,F**). Altogether, our data suggest that NBL cells are sensitive to PLK4i in two distinct ways, depending on the inhibitor concentration: at lower doses, NBL cells are killed by PLK4i regardless of *TRIM37* and *TP53* status, whereas at higher doses the cellular sensitivity depends on high TRIM37 and a functional p53 mitotic surveillance pathway.

### CRISPR screens confirm sensitivity of NBL cells to centrosome overduplication

Our cell viability studies implied that NBL cells are sensitive to low concentrations of PLK4i that lead to centrosome amplification. We therefore next aimed to understand whether it is centrosome amplification *per se* that underlies NBL sensitivity, or another mechanism. We first adopted an unbiased approach using CRISPR/Cas9-enabled screening, to map the genetic networks controlling sensitivity to low doses of RP-1664. This approach has previously proven useful to uncover the mechanism-of-action of various anti-cancer agents, including PLK4i^31,41,42^. To map TRIM37- and p53-independent mechanisms of NBL sensitivity to RP-1664, we transduced CHP134 *TRIM37-low/TP53-KO* cells with the genome-wide TKOv3 Cas9/sgRNA library^43,44^ and 6 days post transduction treated the resulting cell pool with 40nM RP-1664 (dose leading to centrosome amplification) or DMSO as a control for another 12-20 days. Over the course of the screen 40nM RP-1664 resulted in ∼80% loss of viability in TKOv3-transduced CHP134 *TRIM37-low/TP53-KO* cells as compared to DMSO-treated cells. At the end we used next generation sequencing to determine, which sgRNAs were enriched in the final RP-1664-treated cell population over the initial cell pool, as these are likely to cause resistance to the compound (**Figure 3A, Methods**). We identified a number of genes, whose median sgRNA representation increased from the starting population upon RP-1664, but not DMSO treatment (**Figure 3B, Supplementary Table 1**). Analyzing the known functions of these genes revealed striking patterns: Out of 41 genes showing at least 10-fold median sgRNA enrichment, 20 were involved in centrosome biology, 3 were implicated in regulation of apoptosis, and 4 were components of the PIDDosome complex (**Figure 3B**). This data provides multiple lines of evidence that it is indeed centrosome overduplication that is responsible for cell death of CHP134 *TRIM37-low/TP53-KO* cells upon low-dose RP-1664 treatment. The presence of multiple structural components of the centrosome (e.g. *STIL, CEP120, CEP152* and several others), as well as *PLK4* itself, in the list of hits suggests that blunting the cell’s ability to produce centrosomes alleviates the cytotoxicity of 40nM RP-1664. Furthermore, the PIDDosome is known to trigger apoptosis upon recognising supernumerary centrosomes^45,46^ and therefore its inactivation likely allows CHP134 cells to survive despite centrosome amplification. The PIDDosome component caspase 2 is known to induce apoptosis through the pro-apoptotic BCL-2 family of factors including BID and BAX^47^, both of which scored as hits in our screen (**Figure 3B**). In aggregate, this data implicates centrosome amplification as the cause of TRIM37- and p53-independent cell death in CHP134 NBL cells.

**Figure 3.**
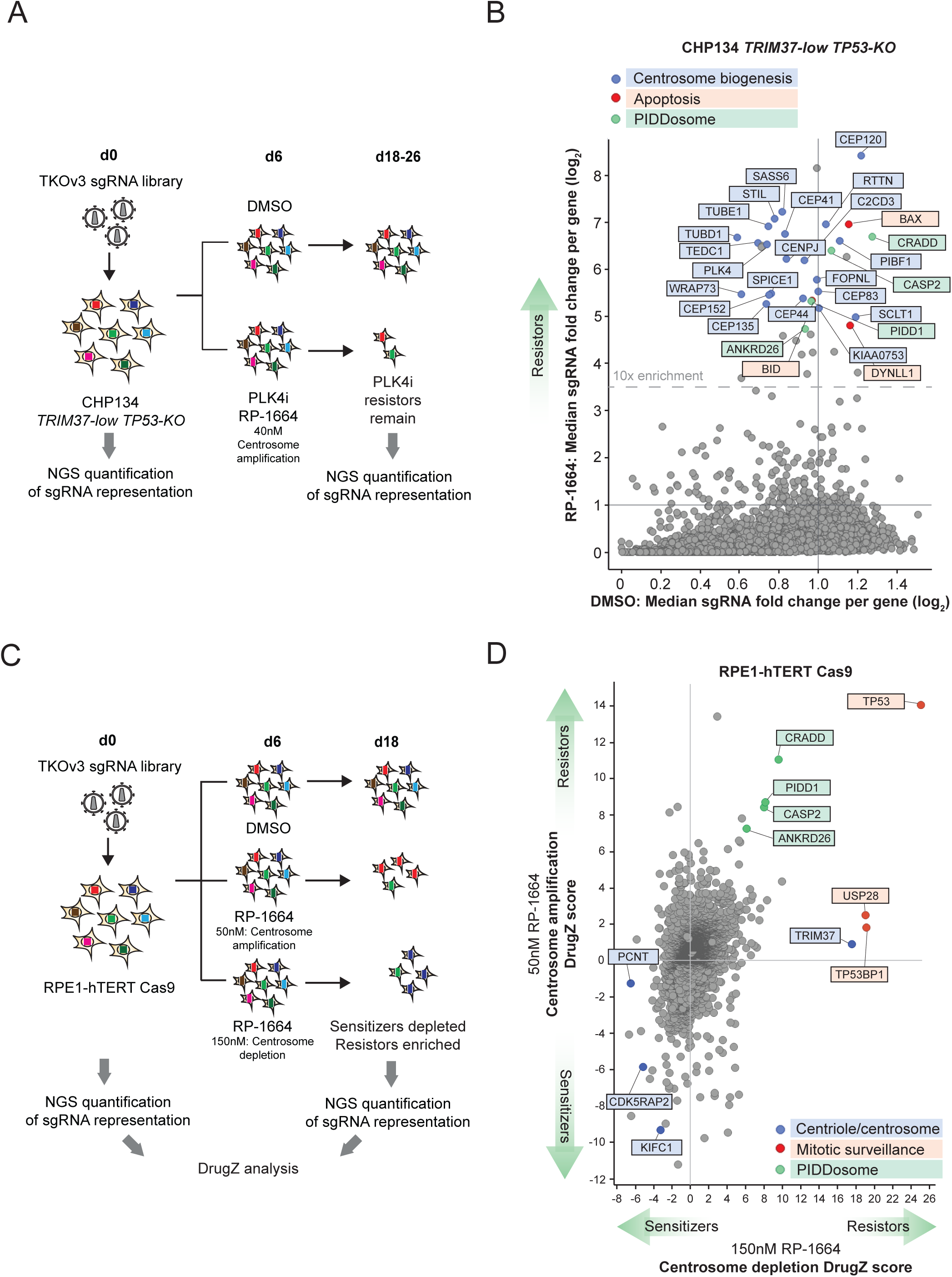
CRISPR screens for genes modulating sensitivity to RP-1664. A,B. Screen for gene knockouts causing RP-1664 resistance in CHP134 *TRIM37-low/TP53-KO* cells. A. Experimental design. See methods for details. B. Screen results. Median sgRNA fold changes per gene in RP-1664-treated cells (Y axis) vs. untreated (X axis). Resistor hits with known functions in centriole/centrosome biogenesis (blue), apoptosis (red) or the PIDDosome (green) are highlighted. C,D. Screen for genes knockouts causing RP-1664 resistance or sensitivity in RPE1-hTERT Cas9 cells. C. Experimental design. See methods for details. D. Screen results. Gene-level DrugZ^70^ scores in cells treated with 50 nM RP-1664 (centrosome amplification; Y axis) vs. 150 nM (centrosome depletion; X axis). Hits with known functions in centriole/centrosome biogenesis (blue), the mitotic surveillance pathway (red) or the PIDDosome (green) are highlighted.

To further explore the genetic framework of response to RP-1664 at doses causing centrosome amplification and doses causing centrosome loss, we performed a second CRISPR/Cas9 chemogenomic screen in Cas9-expressing RPE1-hTERT *TP53-WT* cells treated with DMSO, 50nM RP-1664 (centrosome amplification), or 150nM RP-1664 (centrosome depletion) (**Figure 3C,D; Supplementary Table 2**). Due to the lower sensitivity of RPE1 cells to RP-1664 compared to CHP134, this approach allowed us to determine not only which gene knockouts lead to RP-1664 resistance, but also which genes, when inactivated, increase RP-1664 sensitivity. The results (**Supplementary Table 2**) were in agreement with our CHP134 *TRIM37-low/TP53-KO* screen and consistent with a similar published screen performed using Centrinone B^31^. The screen successfully identified *TRIM37*, and the mitotic surveillance factors *USP28* and *TP53BP1,* as required for cell sensitivity to the dose of RP-1664 leading to centrosome depletion. At the same time, PIDDosome components were among the strongest resistor hits at the lower concentration that leads to centrosome amplification (**Figure 3D**; note that the PIDDosome factors scored as hits also in the higher dose arm, albeit not as strongly as *TRIM37* or *TP53BP1*). These data support the interpretation that cells can become sensitive to RP-1664 in two complementary mechanisms - centrosome depletion and amplification.

In our RPE1 screen, inactivation of *TP53* caused resistance to both doses of RP-1664 (**Figure 3D**). This was confirmed in another screen in RPE1 cells, which we engineered for cytosine-to-adenine CRISPR base-editing by expression of the CBE^FNLS^ dCas9-APOBEC cytidine deaminase fusion^48^ (**Supplementary Figure 4A**). We adapted a previously published sgRNA library capable of generating 5855 single-nucleotide variants (SNVs) in 298 cancer-related genes^49^ to determine which of these SNVs modulate sensitivity to doses of RP-1664 causing centrosome amplification and depletion (50nM and 150nM, respectively). sgRNAs introducing mutations in *TP53* caused resistance to both doses of RP-1664 (with a modest preference for the 150nM dose leading to centrosome depletion; **Supplementary Figure 4B,C; Supplementary Table 3**). This suggests that p53 can be involved in the cellular response to PLK4 inhibition regardless of the underlying mechanism (centrosome amplification or depletion). In contrast, our CHP134 and CHP212 *TP53-KO* data (**Figure 2E,F**) suggest that, whereas active p53 is indispensable for PLK4i sensitivity at doses causing centrosome depletion, cell death at lower concentrations in NBL can occur in absence of p53 activity, potentially highlighting redundancies in cell death activation or cell type-specific differences.

### NBL cells are sensitive to low dose RP-1664 due to inability to tolerate excess centrosomes

In our genome-wide CRISPR/Cas9 RPE1 screen, one of the top genes whose inactivation sensitized to 50nM RP-1664 was *KIFC1* (**Figure 3D**). Its gene product, KIFC1 (also known as HSET) is a kinesin family motor protein previously implicated in cellular tolerance to supernumerary centrosomes^50,51^. KIFC1 facilitates centrosome clustering if more than two centrosomes are present, so that the cell can form a (pseudo-)bipolar spindle and divide normally^51^. This suggested that the inability to cluster supernumerary centrosomes may be a mechanism of sensitivity to low concentrations of RP-1664.

To test whether the lack of centrosome clustering is the basis of NBL sensitivity to low doses of RP-1664, we employed live cell imaging. We stained DNA and microtubules with specific fluorescent dyes (SPY650-DNA and SPY555-Tubulin) and followed mitotic progression of CHP134 NBL cells with or without 50nM RP-1664 treatment. As a ‘normal’ cell control we used RPE1-hTERT *TP53-WT* cells. Whereas under unperturbed conditions CHP134 cells divided normally, we observed multipolar segregation once cells were treated with 50nM RP-1664 (**Figure 4A; Supplementary Video 1,2**). In contrast, RPE1-hTERT cells displayed a wider range of phenotypes upon RP-1664 treatment: although we did observe multipolar segregation in some cases, RPE1 cells were also able to perform pseudo-bipolar mitoses when multiple centrosomes were present, either by centrosome clustering or exclusion of excess centrosomes (**Figure 4A; Supplementary Video 3-5**) – both known mechanisms of adaptation to supernumerary centrosomes^52–54^. To better quantify the frequency of multipolar and (pseudo-)bipolar mitoses in NBL versus normal immortalized cells, we fixed and stained with anti-γ-Tubulin and H3-pS10 antibodies three NBL cell lines (CHP134, CHP212, SHSY5Y), as well as RPE1 *TP53-WT* and *TP53-KO* controls, with or without 50nM RP-1664 treatment. We then manually counted the frequency of cells in anaphase or telophase that were undergoing bipolar versus multipolar segregation (**Figure 4B**). Whereas normal immortalized RPE1 cells (regardless of *TP53* status) only infrequently underwent multipolar mitosis in presence of 50nM RP-1664, multipolar mitoses were common in NBL cells, suggesting that it is indeed the inability to cope with supernumerary centrosomes (by their clustering or inactivation) that underlies PLK4i hypersensitivity in NBL (**Figure 4B**). To examine whether disrupting centrosome clustering would sensitize the normally resistant RPE1 cells to low dose PLK4i (and to validate our prior CRISPR screen result), we inactivated *KIFC1* in RPE1-hTERT Cas9 *TP53-KO* cells using CRISPR/Cas9 and confirmed that the frequency of multipolar cell divisions robustly increased in two resulting *KIFC1-KO* clones upon 50nM RP-1664 treatment (**Figure 4C-E**). Consistent with our CRISPR screen data, *KIFC1-KO* cells were more sensitive to doses of RP-1664 causing centrosome amplification than parental cells, whereas sensitivity to doses leading to centrosome depletion was similar between parental and *KIFC1-KO* cell lines (**Figure 4F**). We conclude that the ability to cluster (or inactivate) supernumerary centrosomes governs cellular sensitivity to partial PLK4 inhibition.

**Figure 4.**
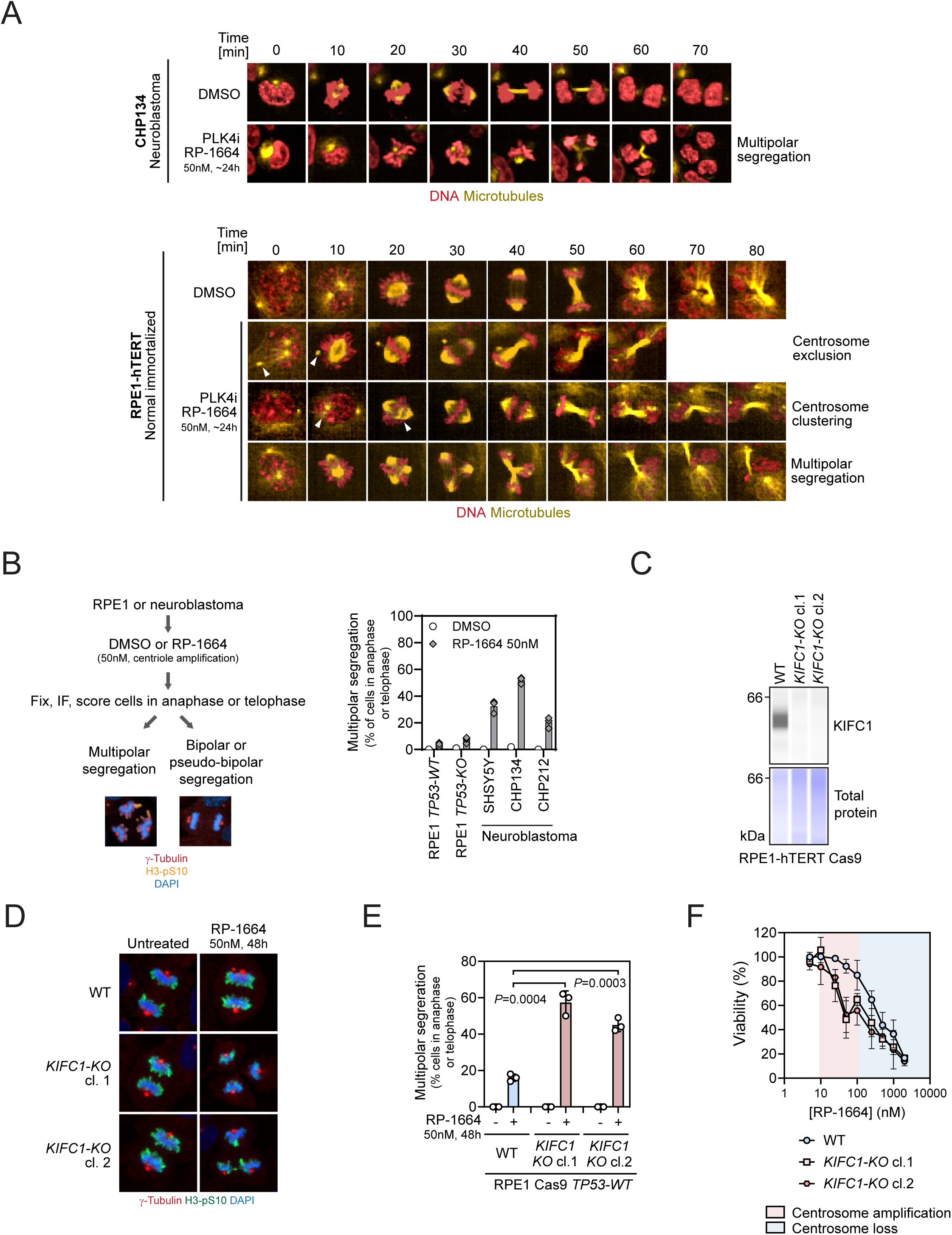
Lack of clustering/exclusion of extra centrosomes leads to RP-1664 sensitivity. A. Representative tempograms from time-lapse imaging of CHP134 (top) and RPE1-hTERT Cas9 (bottom) cells stained with SPY555-Tubulin for microtubules (yellow) and SPY650-DNA for DNA (red) and treated with DMSO or 50nM RP-1664. Example cells undergoing normal division, multipolar segregation, or pseudo-bipolar division with centrosome clustering or centrosome exclusion are shown. B. Left: Workflow for quantification of multipolar segregation frequency. RPE1 or NBL cells were treated with 50nM RP-1664, fixed and immunostained for centrosomes (γ-Tubulin) and mitosis (H3-pS10). DAPI was a nuclear counterstain. Anaphase and telophase cells undergoing either bipolar or multipolar division were quantified. Right: Frequency of multipolar segregation in anaphase or telophase with or without RP-1664 in indicated cell lines. Data from *N*=3 independent experiments (open symbols) with mean (bars) ±SD. C. Representative KIFC1 capillary immunodetection of *KIFC1-WT* and *KIFC1-KO* RPE1-hTERT Cas9 cells. Total protein is a loading control. D. Representative micrographs of *KIFC1-WT* and *KO* cells processed for immunofluorescence with γ-Tubulin and H3-pS10 antibodies with or without RP-1664 treatment. DAPI used as a nuclear counterstain. E. Quantification of *KIFC1-WT* and *KO* cells in anaphase or telophase undergoing multipolar vs. bipolar division in presence or absence of RP-1664. Data from *N*=3 independent experiments (open symbols) with mean (bars) ±SD. *P* values determined with an unpaired two-tailed T-test. F. RP-1664 sensitivity of *KIFC1-WT* and *KIFC1-KO* cells. Mean viability measurements from *N*=3 independent Incucyte growth assays ±SD.

### *TRIM37-* and *TP53-*independent sensitivity of NBL tumors to RP-1664 *in vivo*

To confirm that PLK4i sensitivity of NBL tumor cells due to centrosome amplification is a relevant mechanism *in vivo*, we engrafted into immunodeficient mice parental CHP134 cells, alongside with a *TRIM37-KO* clone (another one than the cell line used in our prior *in vitro* studies but showing the same RP-1664 sensitivity pattern - **Supplementary Figure 5A**) and a *TP53-KO* clone. We confirmed that the resulting tumors maintained the desired TRIM37 and p53 status (**Supplementary Figure 5B**) and treated the mice with either blank chow or a 300PPM RP-1664 chow dose, which throughout each day of dosing led to a free plasma RP-1664 concentration of at or below 100nM on average (**Supplementary Figure 5C**) – i.e. a dose which is *in vitro* associated with centrosome amplification. This regimen led to robust efficacy of RP-1664 in the parental CHP134 model, with tumor regression in 6/7 tumors (**Figure 5A**). Interestingly, inactivation of neither *TRIM37* nor *TP53* affected the efficacy of RP-1664 (**Figure 5A**), suggesting that the TRIM37 and p53-independent mechanism of sensitivity in CHP134 cells is active *in vivo*. The treatment was well tolerated in all models with no BW loss (**Figure 5B**). Altogether, our data suggests that centrosome amplification by low doses of RP-1664 is sufficient to elicit robust anti-tumor activity in an NBL model.

**Figure 5.**
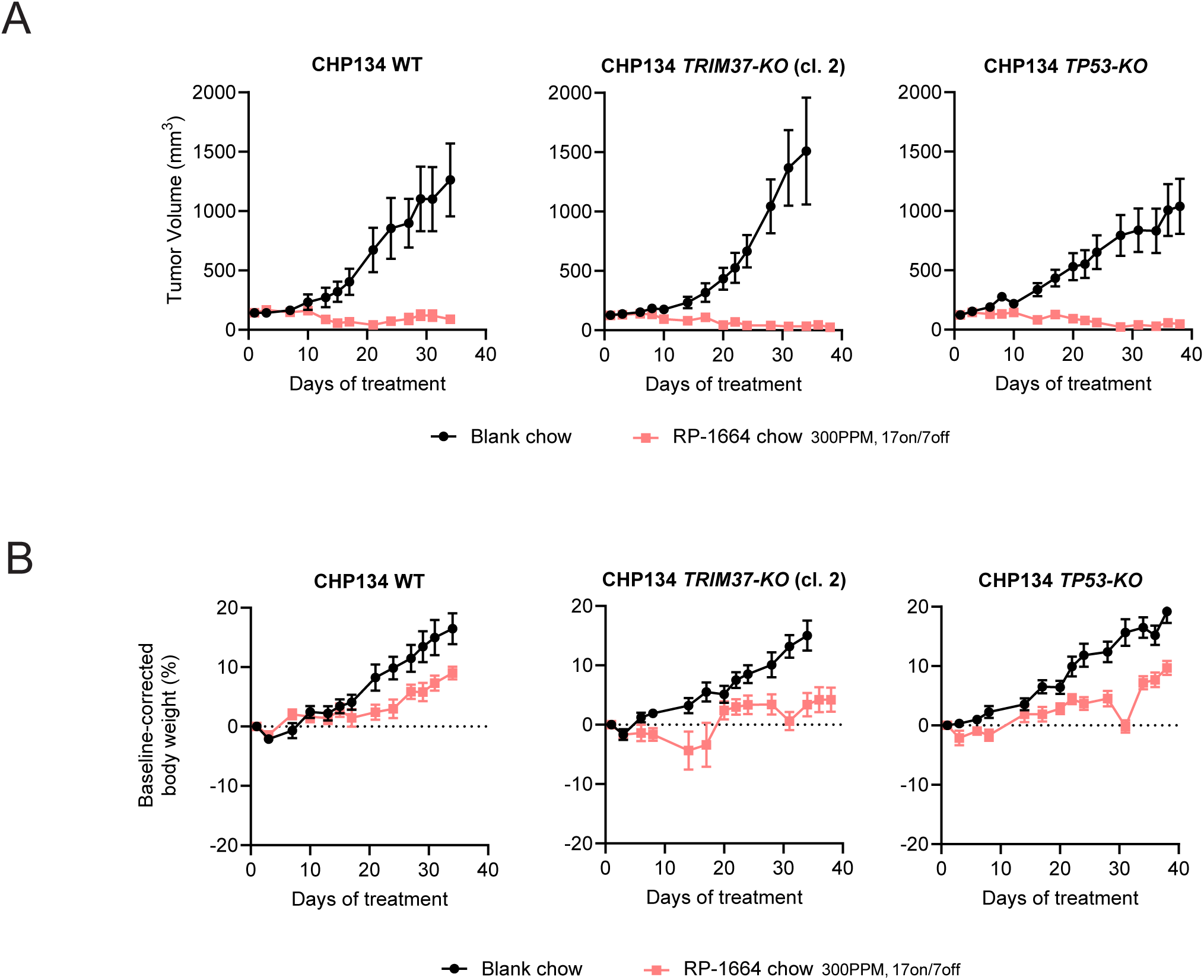
TRIM37- and p53-independent PLK4i sensitivity of NBL tumors *in vivo*. A. Tumor volume measurement of CHP134 WT, *TRIM37-KO* and *TP53-KO* mouse xenografts treated with blank chow or 300ppm RP-1664 chow using a 17d on / 7d off schedule. B. Baseline-corrected body weight change in mice bearing *CHP134 WT*, *TRIM37-KO* and *TP53-KO* xenograft tumors upon indicated treatments. Values in A,B are mean of 7 mice/group ±SEM.

## DISCUSSION

PLK4 inhibition has been suggested as a potential therapeutic strategy for high-risk NBL, based on high prevalence of *TRIM37* gain in this patient population^14^. We now show that PLK4i sensitivity of NBL cells is in fact a composite of two complementary but distinct mechanisms. At lower doses, PLK4i (including RP-1664) amplify centrosomes due to stabilization of partially active PLK4^31–36^ . NBL cells are unable to tolerate supernumerary centrosomes, as they do not employ centrosome clustering or exclusion. In turn NBL cells undergo multipolar segregation, presumably leading to aneuploidy and genomic instability (**Figure 6**). On the other hand, at higher PLK4i concentrations NBL cells lose centrosomes and succumb to mitotic delay or failure as a consequence of high TRIM37 levels^14^ (**Figure 6**). This dual mechanism of sensitivity makes NBL an attractive target population for PLK4i like RP-1664.

**Figure 6.**
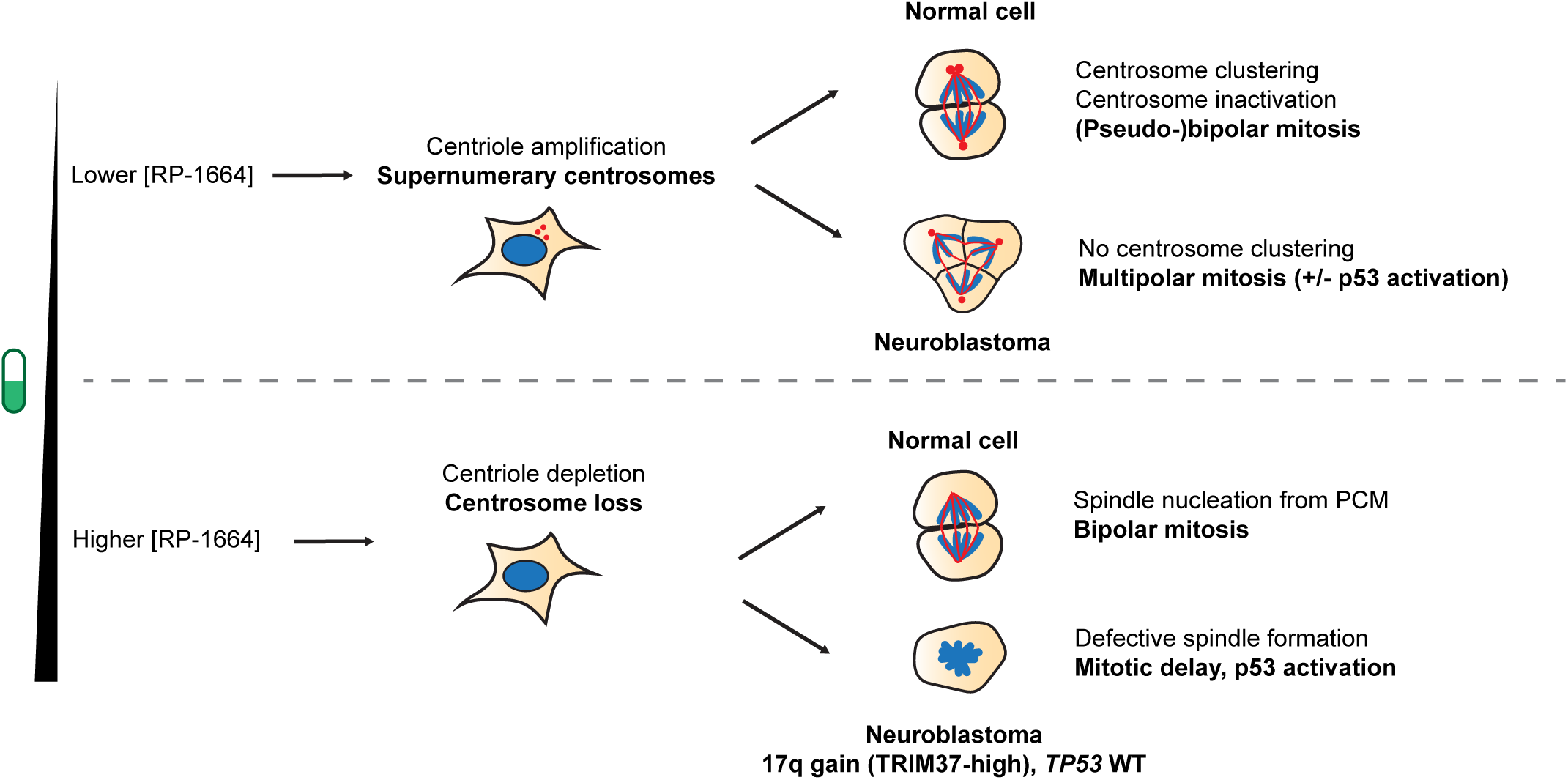
A two-component model of NBL sensitivity to PLK4i. See main text for details.

We demonstrate that centrosome clustering is a *bona fide* mechanism of resistance to PLK4i, as inactivation of a centrosome clustering factor, *KIFC1*, sensitizes normally resistant cells to low dose RP-1664. However, an important outstanding question is what exact genetic basis underlies the inability to compensate for supernumerary centrosomes in NBL. In addition to *KIFC1*, multiple genetic pathways have been previously described to facilitate centrosome clustering^52^, including factors involved in microtubule organization, the spindle assembly checkpoint, sister chromatid cohesion, as well as the actin cytoskeleton and cell adhesion^51,53,55–59^. However, it is unclear whether defects in any of these mechanisms contribute to NBL malignancy. Similarly, it will be important to determine whether any of the recurrent genetic alterations in NBL such as *MYCN* and/or *ALK* gain-of-function, *LIN28B* overexpression, and 1p or 11q loss^9^, contribute to defects in centrosome clustering. Multiple potential links between these alterations and centrosome biology exist. For example, *LIN28B* overexpression has been linked to overactivation of the mitotic kinase Aurora A (AURKA)^60^ involved in centrosome maturation, spindle assembly and cytokinesis^61^. In some studies (although not others^62^), *MYCN* amplification has been associated with centrosome amplification^63,64^. Future correlative and functional studies are thus warranted to identify biomarkers predictive of sensitivity to low-dose PLK4i in individual NBL tumors.

Another important area of investigation is to identify whether the inability to cluster supernumerary centrosomes determines tumor sensitivity to PLK4i outside NBL. In a recent pre-print, Moreno-Marin *et al.* describe a remarkable diversity in centrosome clustering proficiency among the NCI60 cancer cell line panel^65^. The authors show that clustering of supernumerary centrosomes is frequent in mesenchymal tumor cell lines as well as leukemias and correlates with epithelial-to-mesenchymal transition (EMT) in breast cancer models^65^. They propose that lineage determination may contribute to centrosome clustering ability. However, as NBL is a tumor of mesenchymal origin and we show that NBL cell lines are in fact devoid of centrosome clustering, it is unlikely that the same mechanism governs centrosome clustering in all tumor cell lines. In any case, it will be important in future studies to correlate sensitivity to low-dose PLK4i and centrosome clustering ability among a broad range of cellular models.

In addition to describing the mechanism of sensitivity in NBL, our study brings additional support for a complex role of the p53 pathway in the response to abnormal centrosome numbers. Whereas in NBL cells p53 was dispensable for cell death at doses of RP-1664 causing centrosome amplification, its inactivation rescued cell sensitivity at concentrations leading to centrosome loss.

On the other hand, our CRISPR/Cas9 screens in RPE1 cells implicate p53 in cellular sensitivity to both lower and higher doses of PLK4i, consistent with published data^31^. The simplest reconciliation of this apparent discrepancy is that cell death upon centrosome amplification can occur both in a p53-dependent as well as independent fashion. Our CRISPR screens implicate the PIDDosome in mediating cell death at centrosome amplification-inducing doses of RP-1664. Canonically, the PIDDosome is activated upon centrosome amplification by recruitment to distal appendages of centrioles by the ANKRD26 protein, leading to caspase 2-mediated cleavage of the p53 inhibitor MDM2, p21 upregulation and cell cycle arrest / death^66^. Consistent with this model, we observed p21 expression in NBL cells at PLK4i concentrations leading to centrosome amplification. However, these concentrations still induced robust growth inhibition even in p53-deficient cells. Caspase 2 is capable of triggering apoptosis in a p53-independent manner, through release of pro-apoptotic factors from mitochondria^67^ and it is therefore possible that this pathway contributes to PLK4i cytotoxicity in NBL when p53 is inactivated. This possibility is supported by our CHP134 CRISPR screen, where inactivation of mitochondrial BCL2-family pro-apoptotic factors BID and BAX led to RP-1664 resistance. Irrespective of the mechanism, our observation that low doses of PLK4i induce cell death even in absence of p53 activity is important, as it suggests that p53 loss may not be a prominent mechanism of acquired resistance to PLK4i in tumors.

Finally, we show that low-dose PLK4i sensitivity of NBL cells is relevant for *in vivo* efficacy. It is not unreasonable to imagine that in animal models as well as in human both mechanisms of sensitivity may operate in parallel, depending on the dose, schedule, pharmacokinetic profile of a given inhibitor and local concentration of PLK4i in each tumor area. Monitoring centrosome numbers and/or the emergence of multipolar mitoses could therefore be useful pharmacodynamic readouts during PLK4i clinical dose optimization.

In conclusion, we propose that inability to cluster or exclude supernumerary centrosomes is a novel factor predictive of PLK4 inhibition sensitivity, in addition to other published biomarkers including high *TRIM37* levels^13,14^ or hyperactivity of β-Catenin signaling^68^. We believe that our results impact the clinical development of PLK4 in an important way – due to the sensitivity to both low and higher concentrations of PLK4i in NBL tumor cells, lower doses may be sufficient to achieve efficacy in NBL than in tumors showing *TRIM37* gain/amplification alone. Our pre-clinical data warrant testing this hypothesis in clinical trials.

## ACKNOWLEDGEMENTS

This study was funded by Repare Therapeutics. We thank Daniel Durocher, J. Ross Chapman and John M. Maris for critical reading of the manuscript and/or important discussions, and François Denis for technical assistance.

## METHODS

### Cell culture

Cell lines were purchased from the following vendors: RPE1, MCF7, CHP212, SHSY5Y, SKNAS, SKNDZ, SKNFI, SKNSH - ATCC; KELLY – Sigma Aldrich; CHP134, IMR32 – DSMZ. RPE1-hTERT Cas9 *TP53-WT* and *TP53-KO* were described before^37,41^. Cells were cultured in the following media (all supplemented with 10% fetal bovine serum (VWR, 76419-584), 100 U/ml penicillin and 100 μg/ml streptomycin (Corning, 30-001-CI)): RPE1, CHP212, CHP134, SKNDZ, SKNFI – DMEM (Corning, 10-013-CV); IMR32, SKNAS, KELLY – RPMI (Corning, 10-104-CV); MCF7 – EMEM (Corning, 10-009-CV) + 1% non-essential amino acids + 10μg/ml bovine insulin; SKNSH – MEM (Corning 10-010-CV); SHSY5Y – MEM : F12K (1:1; Corning 10-090-CV). All cells were cultured at 37°C and 5% CO_2_.

### CRISPR/Cas9 knockout of *TRIM37*, *TP53, KIFC1*

Cells were transfected with Cas9:sgRNA complexes using Lipofectamine CRISPRmax (Thermo Fisher Scientific) according to manufacturer’s protocol. Cells were allowed to recover for 2-3 days and then seeded for clonogenic outgrowth. Knockout (KO) clones were characterized by Sanger sequencing and ICE analysis^69^ and/or immunoblotting/Simple Western analysis. The following sgRNA target sequences were used: *TRIM37* – TCGCATCAGTGTGCACTTTG or GATGAAGTAAATCAGCTCGA; *TP53* – CAGAATGCAAGAAGCCCAGA; *KIFC1* – GTCCCCCCTATTGGAAGTAA.

### Chemical compounds

RP-1664 was synthesized in house (synthesis will be described in a separate manuscript). Centrinone B was purchased from MedChem Express. Compounds were made up at stock concentrations of 10 mM from powder in dimethyl sulfoxide (DMSO) and kept at −20°C for long-term storage.

### Simple Western capillary immunodetection

For detection of PLK4 stabilization, 150,000 RPE1 cells/well were plated in 12-well plates (Falcon, 353043) in 1 ml/well of media. The next day, increasing doses of RP-1664 were added to the cells using a Tecan D300E digital dispenser. 48 hours later, cells were washed with 1 ml of PBS and lysed in-plate with 50 µl of RIPA buffer (Sigma, R0278-50ML), supplemented with 1X protease and phosphatase inhibitors (Thermo-Fisher, 78440).

For TRIM37 and p21 immunodetection, cell lysates were prepared in an analogous manner, without RP-1664 treatment.

Xenograft tumor lysates were prepared as follows:

Following lysis, protein concentrations were measured using a DC Protein Assay Kit (Bio-Rad, 5000112) and protein levels in all samples were normalized to 1 µg/µl in lysis buffer. Immunodetection was performed using a Simple Western JESS instrument (Bio-Techne) according to the manufacturer’s instructions. 3 µl of lysate was loaded per capillary and proteins were detected with their respective primary antibodies (see below). Separation time was set to 25 min., separation voltage set to 375V, primary antibody incubation time to 90 min, and secondary antibody incubation time to 60 min.

### Immunoblotting

Cells were lysed directly in 2x Novex Tris-Glycine SDS sample buffer + 200mM DTT (Thermo Fisher Scientific LC2676) at a concentration of 1×10^6^ cells/ml and boiled at 95°C for 5 min. 20-30μl of cell lysate was run on a Novex Tris-Glycine SDS gel (Thermo Fisher Scientific). Proteins were then transferred to a 0.2μM nitrocellulose membrane at 90V for 2-2.5h. Membranes were blocked with 5% milk/TBST for 30 min at room temperature (RT) and incubated with a primary antibody at 4°C overnight. Membranes were then washed 3x 5min with TBST and incubated with a secondary antibody for 1h at RT, after which they were washed again and developed using the Pierce ECL Pico or Femto Western Blotting substrate (Thermo Fisher Scientific). Chemiluminescence was detected using an Amersham Imager 680 instrument (GE).

### Immunofluorescence

Cells were seeded on black, clear bottom, poly-D-lysine coated 96-well imaging plates (PhenoPlate 96; Revvity, 6055500) and increasing concentrations of RP-1664 were added the following day using a Tecan D300E automatic dispenser. Cells were fixed 48-72 h later as follows: Media was removed, cells were rinsed with PBS, and fixed with 4% paraformaldehyde/PBS (PFA; Thermo Fisher, J19943.K2) for 10 mins at room temperature (RT). Plates were then rinsed 2x with PBS and stored at 4°C. For immunofluorescent staining, cells were permeabilized with 0.3% Triton X-100/PBS for 30 min at RT, and rinsed 2x with PBS. Plates were subsequently blocked with blocking buffer (5% normal goat serum / 0.1% Triton X-100 / PBS) for 1 h at RT. Blocking buffer was removed and cells were incubated overnight at 4°C or 2h at RT with primary antibodies in blocking buffer. The next day, plates were rinsed 3x with 0.1% Triton X-100/PBS and incubated for 1h at RT with a secondary antibodies 0.5 μg/ml 4′,6-diamidino-2-phenylindole (DAPI) was included in the secondary antibody dilution as a DNA counterstain. Plates were rinsed 3x with 0.1% Triton X-100/PBS and PBS was added to image on an Operetta CLS or Opera Phenix automated high-content microscope in confocal mode, using a 40x NA1.1 water immersion objective (Revvity).

### Cell growth assays

Cells were seeded on white, clear bottom 96-well plates (Corning, 3903) and compounds were added the next day using a Tecan D300E automatic dispenser. The following seeding densities were used (cells/well): RPE1 – 300; CHP134 – 1500; CHP212 – 1000; SHSY5Y – 1000; IMR32 – 700; SKNFI – 3000; SKNSH – 500; SKNDZ – 2000; KELLY – 700; SKNAS – 700-800. When cells reached near-confluence (∼4-6 population doublings / ∼6-12 days), plates were imaged on an Incucyte S3 automated microscope. Images were analyzed using the Incucyte 2022B software to determine % confluence. Alternatively, cells were processed using the CellTiter Glo cell viability assay (Promega) according to manufacturer’s instructions and luminescence was measured with a FlexStation 3 plate reader. Data were normalized to untreated controls and IC_50_ values were obtained by non-linear least squares fitting of the normalized data to a three-or four-parameter dose-response model in GraphPad PRISM 10.

### Cytotox and Annexin V cell viability assays

Cells were seeded at low density on black poly-lysine coated 96-well plates (Revvity, 6055500). Two days after seeding, cells were incubated with Incucyte dyes to detect cell death (Cytotox; Sartorius 4633; 1:500) or apoptosis (Annexin V; Sartorius 4642; 1:2000). Serial dilutions of RP-1664 were added to the plate using a Tecan D300E digital dispenser. Plates were imaged on an Incucyte S3 automated microscope in both phase-contrast and green fluorescent channels to simultaneously monitor cell confluence and viability. Images were captured every 8 hours for 5 days and analyzed using Incucyte 2022B software. Cell death induction by RP-1664 was analyzed at the time each cell line reached 2 doublings (CHP134: 48h; CHP212: 80h), and was calculated as the % area stained with green dye (Cytotox or Annexin V) normalized to % confluence, and relative to DMSO.

### Antibodies

The following antibodies and dilutions were used for immunoblotting (IB), immunofluorescence (IF) or capillary immunodetection (JESS): rabbit anti-PLK4 E6A7R (Cell Signaling Technologies 71033; JESS 1:200), rabbit anti-TRIM37 (Bethyl A301-174A; JESS 1:50), mouse anti-p53 DO-1 (sc-126; IB 1:1000), rabbit anti-p21 12D1 (Cell Signaling Technologies 2947; IF 1:500, JESS 1:300), rabbit anti-γ-Tubulin EPR16793 (Abcam 179503; IF 1:500), mouse anti-H3pS10 3H10 (Sigma Aldrich 05-806; IF 1:1000), rabbit anti-KIFC1 (ProteinTech 20790-1-AP; JESS 1:50), rabbit anti-DYKDDDDK tag (FLAG) D6W5B (Cell Signaling Technologies 14793; IB 1:1000), rabbit anti-Vinculin E1E9V (Cell Signaling Technologies 13901; IB 1:5000), Alexa Fluor 488/555/647-conjugated goat anti-rabbit or anti-mouse IgG (H+L) (Thermo Fisher Scientific A-11008/A-21428/A-21245/A-11001/A-21422/A-21235; IF 1:500-

1:1000), HRP-conjugated goat anti-mouse IgG (BioRad L005680; 1:5000 IB), HRP-conjugated goat anti-rabbit IgG (Jackson ImmunoResearch 111-035-144; IB 1:5000), HRP-conjugated anti-rabbit secondary antibody (Bio-Techne 042-206; JESS undiluted).

### Live cell imaging

Cells were seeded on 96-well poly-lysine coated PhenoPlates (Revvity, 6055500) and incubated in phenol red-free media supplemented with SPY650-DNA (Cytoskeleton CY-SC501) and SPY555-Tubulin (Cytoskeleton CY-SC203) at 1:6000 dilution for 2 hours before imaging. RP-1664 or DMSO treatments were added 24 hours prior and maintained during imaging. Time-lapse images were captured on an Opera Phenix automated high-content microscope in confocal mode under controlled environmental conditions (37C and 5% CO_2_), using a 40x NA1.1 water immersion objective (Revvity). Images were captured every 4-5 minutes for ∼6 hours and analyzed manually using Harmony. Mitotic duration was recorded for each dividing cell, counted as the time from nuclear envelope breakdown to mitotic exit (i.e., anaphase or chromatin decondensation).

### CRISPR/Cas9 chemogenomic screening

CRISPR/Cas9 screens were performed as previously described^44^. The RP-1664 resistance screen was performed in a CHP134 *TRIM37-low TP53-KO* neuroblastoma cell line. Cells were transduced with lentivirus carrying the TKOv3 sgRNA library at a multiplicity-of-infection (MOI) of ∼0.3. The screen was conducted in technical duplicates, and library coverage of >100 cells per sgRNA was maintained at every step. Puromycin-containing medium (0.5 µg/ml) was added 2 days after infection to select for transductants. Selection was continued until 96 h after infection, which was considered the initial time point (*t*_0_). RP-1664 was added to the cells starting from day 6 (*t*_6_) at 40 nM, corresponding to LD80. From *t*_10_ onwards, RP-1664-containing medium was subsequently refreshed every four days until the screen was terminated at *t*_26_. To identify genes whose deletion caused resistance to RP-1664, genomic DNA was isolated from surviving cells using the QIAamp Blood Maxi Kit (Qiagen) and genome-integrated sgRNA sequences were amplified by PCR using NEBNext Ultra II Q5 Master Mix (New England Biolabs). i5 and i7 multiplexing barcodes were added in a second round of PCR and final gel-purified products were sequenced on an Illumina NextSeq500 system to determine sgRNA representation in each sample.

A second CRISPR/Cas9 chemogenomic screen in Cas9-expressing RPE1-hTERT *TP53-WT* cells treated with DMSO, 50nM RP-1664 (centrosome amplification), or 150nM RP-1664 (centrosome depletion) was performed with cells transduced with the TKOv3 sgRNA library at an MOI of 0.4. Technical duplicates for each condition and library coverage of >400 cells per sgRNA were maintained throughout the screen procedure. 2-days post-infection, transduced cells were selected by adding puromycin to 2 μg/ml in the growth media for 48h. As above, the initial timepoint (*t*_0_) was 96h post-infection. At (*t*_6_), RP-1664 was added to the cells at 50 nM and 150 nM, corresponding to centriole amplification and depletion doses, respectively. Media containing the indicated concentrations of RP-1664 was refreshed every four days until the screen reached 18 days. To identify genes whose deletion caused sensitivity and/or resistance to RP-1664, genomic DNA was isolated from surviving cells at every timepoint, and processed as described above. Sample data analysis was performed using the DrugZ (https://github.com/hart-lab/drugz)^70^ algorithm.

### Base editing screens

RPE1-hTERT *TP53-WT* stably expressing CBE^FNLS 48^ were transduced with lentivirus carrying a sgRNA expression library based on the HBES^49^ (lacking the sensor) at a multiplicity-of-infection (MOI) of ∼0.3. The screen was conducted in technical triplicates, and library coverage of >1,000 cells per sgRNA was maintained at every step. Puromycin-containing medium (2 µg/ml) was added 2 days after infection to select for transductants. Selection was continued until 96 h after infection, which was considered the initial time point (*t*_0_). RP-1664 was added to the cells at day 6 (*t*_6_) and 10 (*t_10_*) at the indicated concentrations and the screen was terminated at *t*_18_. Sample processing was carried out as in CRISPR/Cas9 chemogenomic screening. FASTQ files were aligned to the library reference sequences using Bowtie to generate read counts for each sample replicate at the guide level. Raw read counts from the *t_18_* timepoint of each treatment condition were compared to DMSO control at T18 using DESeq2 to calculate log_2_FoldChange values, and the PCR2 forward primer used for each sample replicate was modeled as a covariate.

### Animal studies

MCF7 mouse xenograft studies were performed at Oncodesign Inc. under regulations from the Canadian Council on Animal Care and the National Research Council Guide. Briefly, female BALB/c Nude mice were supplemented with estradiol in drinking water (2.5 µg/mL) and gamma irradiated (1.2 Gy) 1 week and 24-72 hours prior to inoculation in the right flank using 10 million cells respectively. Animals were placed under blank chow for acclimatization 3 – 5 days prior to randomization. Mice were randomized into 6 groups of 6 animals each when tumor volume reached a mean of 100 – 200 mm^3^ and treatment with RP-1664 formulated chow began thereafter. Body weights and tumor volume were both measured twice a week, the latter being assessed with calipers. Tumor volume was calculated using the formula 0.52×L×W^2^, with percent tumor growth inhibition (% TGI) defined as: % TGI= ((TVvehicle_last_ – TVvehicle_day0_) - (TVtreated_last_ – TVtreated_day0_)) / (TVvehicle_last_ – TVvehicle_day0_) x 100.

CHP134 mouse xenograft studies were performed at Repare Therapeutics in a vivarium accredited by the Canadian Council on Animal Care with an Institutional Animal Care Committee-approved protocol. Briefly, female CB/17 SCID mice were inoculated in the right flank with 10 million cells of either parental CHP134, CHP134 *TP53-KO* or CHP134 *TRIM37-KO* and animals placed under blank chow for acclimatization 3 – 5 days prior to randomization. Mice were grouped according to their tumor phenotype and randomized into 3 groups of 7 animals per cell line when tumor volume reached a mean of 100 – 150 mm^3^; treatment with RP-1664 formulated chow began thereafter. Body weights and tumor volume were both measured three times a week, the latter being assessed with digital calipers. Tumor volume was calculated using the formula 0.52×L×W^2^, with percent tumor growth inhibition (% TGI) defined as: % TGI= ((TVvehicle_last_ – TVvehicle_day0_) - (TVtreated_last_ – TVtreated_day0_)) / (TVvehicle_last_ – TVvehicle_day0_) x100. Percentage changes in body weight (% BW) was calculated based on individual body weight changes relative to the start of treatment, using the formula: % BW change = (BW_last_-BW_day0_/ BW_day0_) x 100.

## Author Contributions

ISB, MCS, MZ: Conceptualization, experimental data collection and analysis, paper writing. EG, LL, CC, AS, HK, DH, NL, JB, AAQ, FV: Experimental data collection and/or analysis. AV, JTFY, MLH, SJM, CGM: Supervision, conceptualization.

---

**Supplementary Figure 1. Related to Figure 1.**
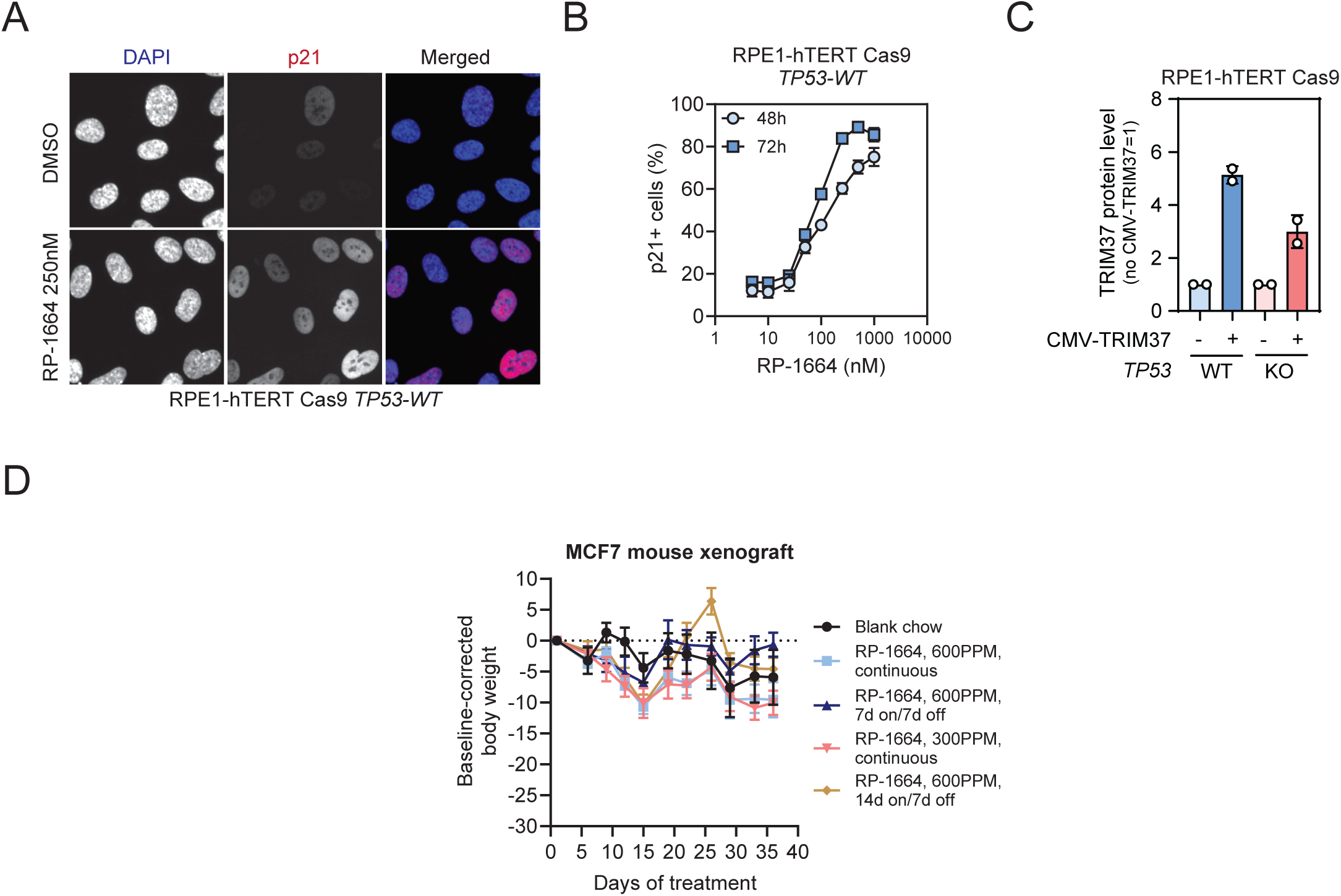
A. Representative micrographs of RPE1-hTERT Cas9 cells after no treatment or treatment with RP-1664 and p21 immunostaining. DAPI is a nuclear counterstain. B. Quantification of p21-positive (p21+) RPE1-hTERT Cas9 cells after 48 or 72h treatment with indicated concentrations of RP-1664. Mean of *N*=3 independent experiments ±SD. C. Quantification of TRIM37 protein levels in RPE1 TP53-WT and KO cells with or without CMV-TRIM37 overexpression. Measured by capillary immunodetection, data from *N*=2 independent experiments (circles) with mean (bars) ±SD. D. Baseline-corrected body weight change in mice bearing MCF7 xenograft tumors upon indicated treatments and schedules. Mean of *N*=6 mice/group ±SEM.

**Supplementary Figure 2. Related to Figure 2.**
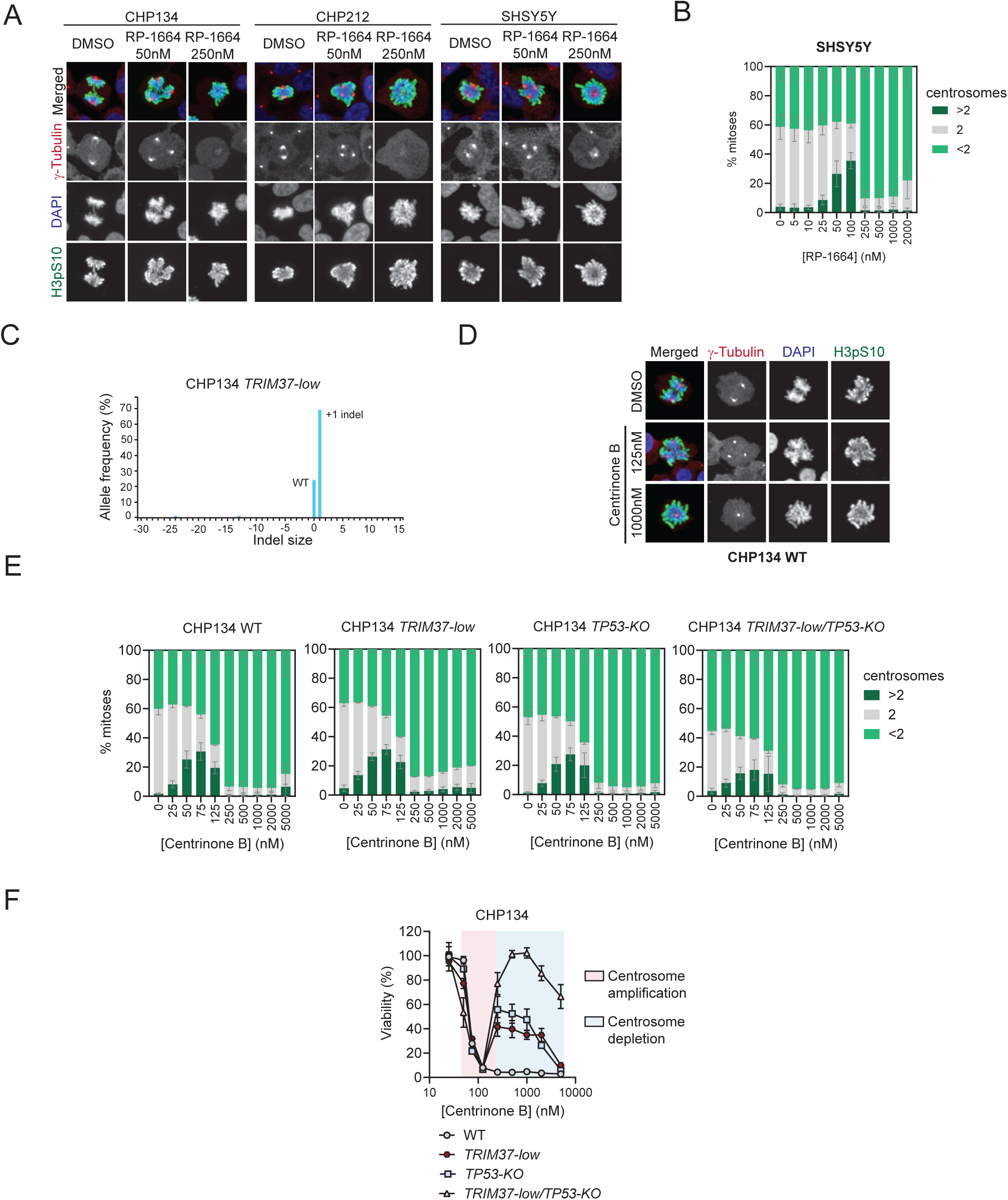
A. Representative micrographs of CHP134, CHP212 and SHSY5Y cells after no treatment of treatment with indicated RP-1664 concentrations and immunofluorescence staining with γ-Tubulin (visualizing centrosomes) and H3-pS10 (mitotic marker) antibodies. DAPI is a nuclear counterstain. B. Quantification of mitotic SHSY5Y cells with <2, 2, and >2 centrosomes at indicated RP-1664 concentrations in *N*=3 independent experiments. Mean value (bars) is shown ±SD. C. ICE^69^ quantification of indel allele frequency in CHP134 *TRIM37-low* cells. D. Representative micrographs of CHP134, CHP212 and SHSY5Y cells after no treatment of treatment with indicated centrinone B concentrations and immunofluorescence staining with γ-Tubulin (visualizing centrosomes) and H3-pS10 (mitotic marker) antibodies. DAPI is a nuclear counterstain. E. Quantification of mitotic CHP134 WT, *TRIM37-low*, *TP53-KO* and *TRIM37-low/TP53-KO* cells with <2, 2, and >2 centrosomes at indicated centrinone B concentrations in *N*=3 independent experiments. Mean value (bars) is shown ±SD. F. Viability of CHP134 cells of indicated genotypes upon treatment with indicated concentrations of centrinone B as measured by Incucyte growth assays. Concentrations inducing centrosome amplification (pink) and depletion (blue) are highlighted. Mean of *N*=3 independent experiments ±SD.

**Supplementary Figure 3. Related to Figure 2.**
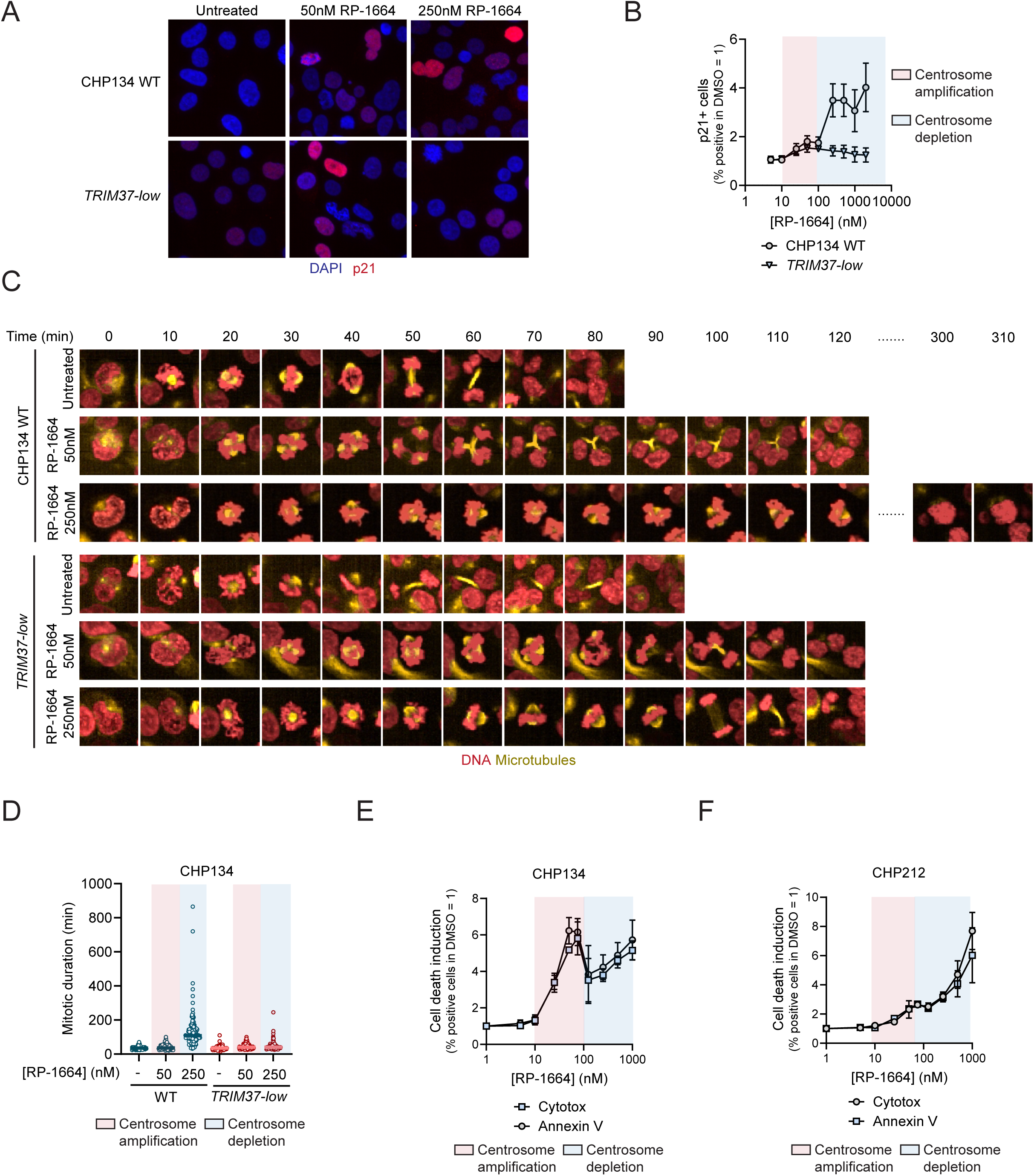
A. Representative micrographs of CHP134 WT and *TRIM37-low* cells after no treatment or treatment with RP-1664 and p21 immunostaining. DAPI is a nuclear counterstain. B. Quantification of p21-positive (p21+) CHP134 WT and *TRIM37-low* cells after 48h treatment with indicated concentrations of RP-1664. Mean of *N*=3 independent experiments ±SD. Concentrations inducing centrosome amplification (pink) and depletion (blue) are highlighted. C. Representative tempograms from time-lapse imaging of CHP134 WT and *TRIM37-low* cells stained with SPY555-Tubulin for microtubules (yellow) and SPY650-DNA for DNA (red) with or without treatment with indicated RP-1664 concentrations. D. Quantification of mitotic length in CHP134 WT and *TRIM37-low* cells by live-cell imaging. Cells were treated with DMSO, 50nM RP-1664 (centrosome amplification) or 250nM RP-1664 (centrosome depletion). Data from *N*=2 independent experiments. Each data point is a single cell with median (solid line). E,F. Induction of cell death by RP-1664 in CHP134 (E) and CHP212 (F) NBL cells. Fold change from DMSO in the percent of Cytotox- and Annexin V-positive cells after indicated RP-1664 concentrations. Mean of N=3 independent experiments ±SD. Concentrations of RP-1664 inducing centrosome amplification (pink) and depletion (blue) are highlighted.

**Supplementary Figure 4. Related to Figure 3.**
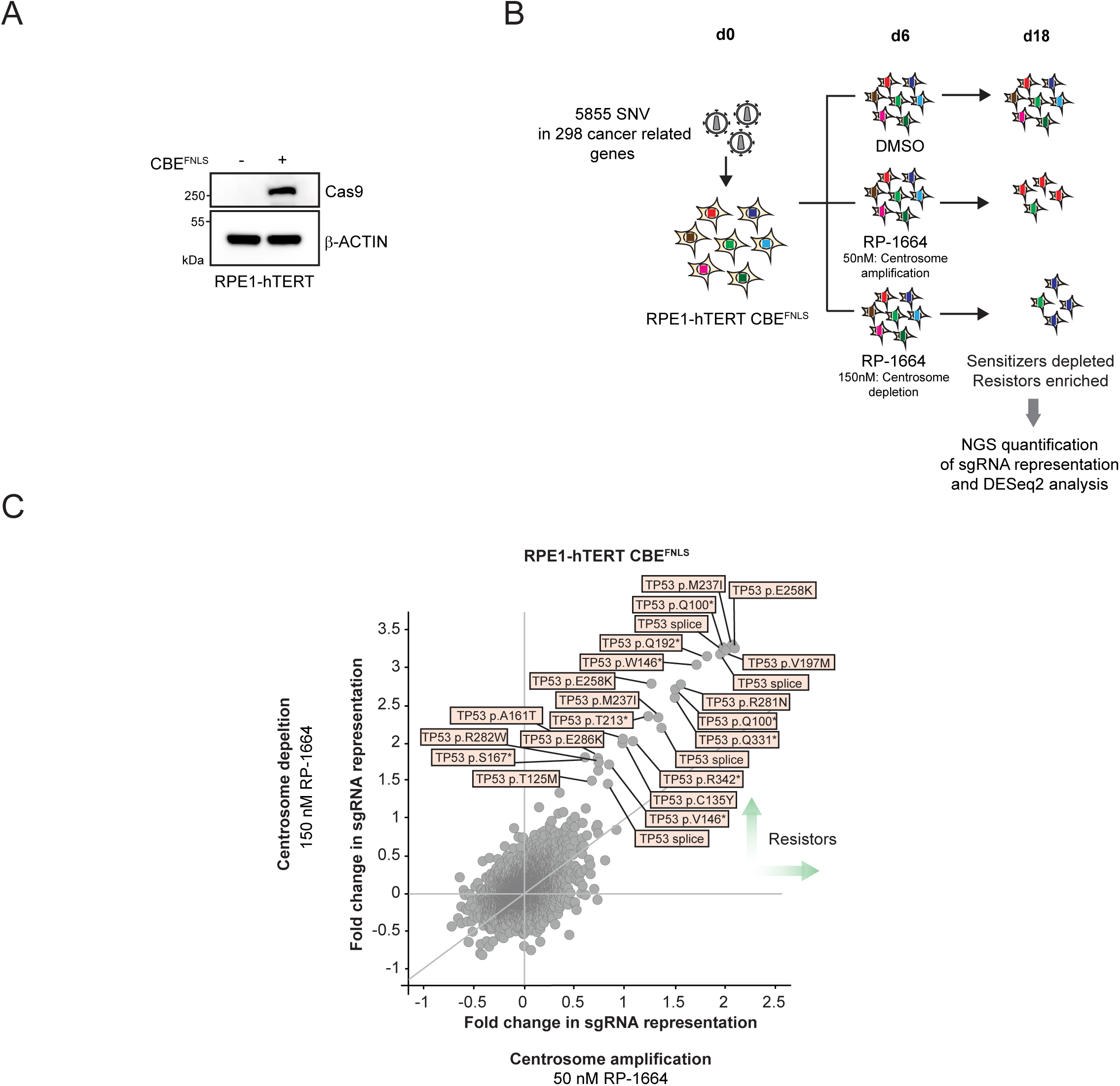
A. Expression of the CBE^FNLS^ base editor in RPE1-hTERT cells. Representative anti-spCas9 immunoblot of whole cell extracts. β-ACTIN is a loading control. B. Experimental design of a base-editing CRISPR screen for cancer-relevant single-nucleotide variants (SNVs) that modulate RP-1664 sensitivity in RPE1-hTERT CBE^FNLS^ cells. See Methods for details. C. Screen results. Fold changes in representation of individual sgRNAs in cells treated with 50nM RP-1664 (centrosome amplification; X axis) vs. 150nM (centrosome loss; Y axis). sgRNAs inducing TP53 mutations that cause RP-1664 resistance are highlighted.

**Supplementary Figure 5. Related to Figure 5.**
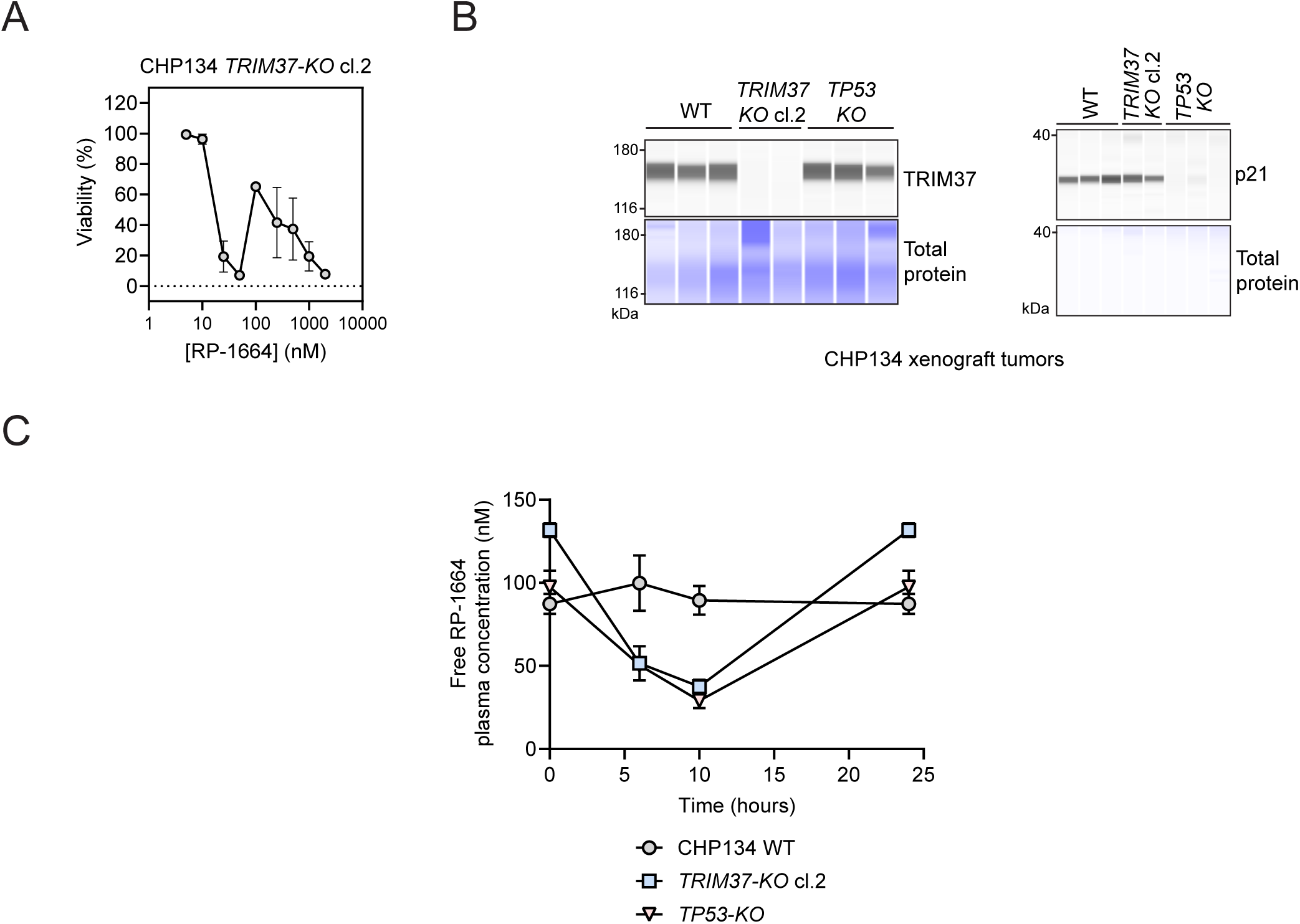
A. RP-1664 sensitivity of CHP134 *TRIM37-KO* clone 2 cells used as a xenograft model. Mean cell viability values from *N*=3 Incucyte growth assays ±SD. B. TRIM37 (left) and p21 (right) capillary immunodetection in tumor lysates of CHP134 xenografts of indicated genotypes. Each lane represents an individual tumor. Total protein is a loading control. C. Free (not bound to plasma protein) plasma concentrations of RP-1664 in mice bearing CHP134 tumors of indicated genotypes treated with 300ppm RP-1664 chow. Mean of *N*=3 mice at indicated time points post dosing is shown ±SEM.

**Supplementary Tables 1-3 will be available at publication**

**Supplementary Video 1,2. Related to Figure 4A**. Example time-lapse video of CHP134 cells with (Supplementary Video 2) or without (Supplementary Video 1) RP-1664 treatment. Microtubules are labeled in yellow, DNA in red.

**Supplementary Video 3-5. Related to Figure 4A**. Example time-lapse video of RPE1-hTERT Cas9 *TP53-WT* cells with (Supplementary Video 4,5) or without (Supplementary Video 3) RP-1664 treatment. Microtubules are labeled in yellow, DNA in red.

